# oCEM: Automatic detection and analysis of overlapping co-expressed gene modules

**DOI:** 10.1101/2021.03.15.435373

**Authors:** Quang-Huy Nguyen, Duc-Hau Le

## Abstract

**Background:** When it comes to the co-expressed gene module detection, its typical challenges consist of overlap between identified modules and local co-expression in a subset of biological samples. The nature of module detection is the use of unsupervised clustering approaches and algorithms. Those methods are advanced undoubtedly, but the selection of a certain clustering method for sample- and gene-clustering tasks is separate, in which the latter task is often more complicated.

**Results:** This study presented an R-package, Overlapping CoExpressed gene Module (oCEM), armed with the decomposition methods to solve the challenges above. We also developed a novel auxiliary statistical approach to select the optimal number of principal components using a permutation procedure. We showed that oCEM outperformed state-of-the-art techniques in the ability to detect biologically relevant modules additionally.

**Conclusions:** oCEM helped non-technical users easily perform complicated statistical analyses and then gain robust results. oCEM and its applications, along with example data, were freely provided at https://github.com/huynguyen250896/oCEM.

## Background

The introduction of genome-wide gene expression profiling technologies observed so far has turned the biological interpretation of large gene expression compendia using module detection methods to be a crucial pillar [1–3]. Here, a module itself is a set of genes that are similarly functioned and jointly expressed. Co-expressed modules do not only help to globally and objectively interpret gene expression data [4, 5], but it is also used to discover regulatory relationships between putative target genes and transcription factors [6–8]. Also, it is useful to study the origin [9] and development [10] of complex diseases caused by many factors.

The nature of module detection is the use of unsupervised clustering approaches and algorithms. Those methods are advanced undoubtedly, but the selection of a certain clustering method for sample- and gene-clustering tasks is separate, in which the latter task is often more complicated. Indeed, users should predetermine the following limitations before applying clustering methods to gene expression. Firstly, not all clustering methods have the ability to tackle the problem of overlap between modules. While clustering patients into biologically distinct subgroups is our ultimate goal, the way to group genes into functional modules need to be more careful since genes often do not work alone; e.g, previous studies have reported that at least five genes work in concert [11] and that their interaction is associated with multiple pathways [12]. Secondly, clustering methods often ignore local co-expression effects which only appear in a subset of all biological samples and instead are interested in co-expression among all samples. This results in loss of meaningful information due to highly context-specific transcriptional regulation [13]. Among existing clustering methods, decomposition methods [14] and biclustering [15] are said to possibly handle the two above restrictions. These obviously affect the selection of which clustering method in the context of gene expression; however, it is rarely examined sufficiently, leading to a typical example is the tool weighted gene co-expression network analysis (WGCNA) [16] with a hierarchical agglomerative clustering [17].

Wouter Saelens *et al* [18] have conducted a holistic comparison of module detection methods for gene expression data and realized that the decomposition methods, including independent component analysis (ICA) [19–21], principal component analysis (PCA) [22], and independent principal component analysis (IPCA) [23], are the best. In this study, we have proposed an R tool, named Overlapping CoExpressed gene Module (oCEM), which integrated these methods in the hope that it could be a potential alternative to rectify the limitations above. In particular, we developed a state-of-the-art statistical method, called *optimizeCOM*, to specify the optimal number of principal components in advance required by the decomposition methods. Then, the function *overlapCEM* available in oCEM supported users to implement the module detection and analysis automatically. These helped non-technical users to easily perform complicated statistical analyses and gain robust results in a surprisingly rapid way. We also demonstrated better performance of oCEM in comparison to other high-tech methods in terms of identification of clinically relevant co-expressed modules.

## Implementation

### Overview of oCEM

Figure 1 shows the automatic framework for module detection and analysis included in oCEM. Gene expression matrix first suffered from the two pre-processing steps: excluding outlier individuals and normalization prior to being the input of oCEM. The result of normalization was that the distribution of each gene expression was centered and standardized across samples. The user now put the data to oCEM, and it printed automatically out the following results: (i) coexpressed gene modules (the module was determined from a particular component by using one of the optional postprocessing steps described above), (ii) hub-genes specific to each module, and (iii) analysis result of associations between each module and each clinical feature of choice (e.g., tumor stages, glycemic index, weight,…). Note that oCEM decomposed the expression matrix into the product of two or more sub-matrices by only using one of the two decomposition methods, including ICA (the *fastICA* algorithm) and IPCA (the *ipca* algorithm). oCEM did not include PCA because of the following reasons: (i) PCA assumes that gene expression follows a Gaussian distribution; however, many recent studies have demonstrated that microarray gene expression measurements follow a non-Gaussian distribution instead [24–27], (ii) The idea behind PCA is to decompose a big matrix into the product of several sub-matrices and then retain the first few components which have the maximum amount of variance. Mathematically, this helps to do dimension reduction, noise reduction, but the highest variance may be inappropriate to the biological reality [28, 29].

**Figure 1.**
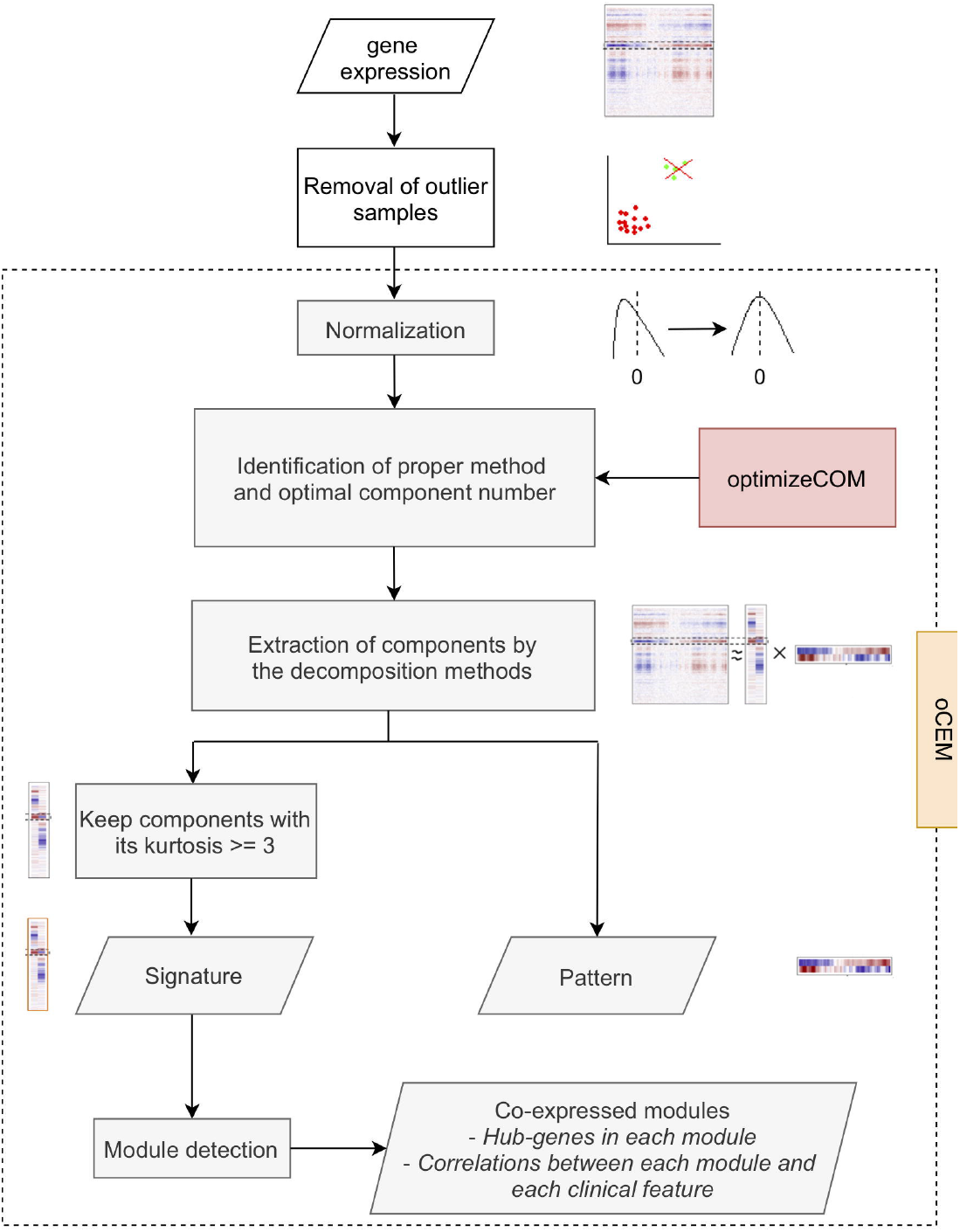
Automatic analysis framework of oCEM. Gene expression data first underwent the two pre-processing steps: removal of outlier samples and normalization. Then, the user could refer to the recommendation of oCEM regarding which decomposition method should be selected and how many component numbers were optimal by using the function *optimizeCOM*. Next, the processed data were inputted into the function *overlapCEM*, rendering coexpressed gene modules (i.e., Signatures with their own kurtosis ≥ 3) and Patterns. Kurtosis statistically describes the “tailedness” of the distribution relative to a normal distribution. Finally, corresponding hub-genes in each module and the association between each module and each clinical feature of choice were identified.

Since the output of the decomposition methods generally consisted of two parts, the components for genes and the components for samples, we, in this study, distinguished the two by the term of signatures and patterns, respectively (Additional File 1:Supplementary Methods and Additional File 1:FigureS1). When it comes to the first matrix product (vertical rectangle in Additional File 1: FigureS1), oCEM described the characteristic of different signatures by, between them, a set of genes of which the overlap was allowed. In contrast, for the second matrix product (horizontal rectangle in Additional File 1:FigureS1), oCEM characterized each component by its expression patterns in biological samples.

#### optimizeCOM algorithm

The first step of oCEM involved deciding how many principal components should be. To support the user to possibly make a good decision, we developed an R function called *optimizeCOM*. The idea behind this function was based on random permutations adapted from [30], aimed not only to help the user to know which method should be selected but also to specify the optimal number of principal components to extract by ICA or IPCA (detailed in Additional File 1:Supplementary Methods and Additional File 1:FigureS2).

#### Keep components with non-Gaussian distribution

oCEM equipped with ICA and IPCA required the distributions of the signatures across genes must be as non-Gaussian as possible; ideally, they should be heavy-tailed normal distributions. Due to this requirement, the kurtosis was recruited, which statistically describes the “tailedness” of the distribution [20], and only kept signatures whose kurtosis value ≥ 3.

#### Detection of co-expressed gene modules

It was evident that a few genes at the tails of a heavy-tailed distribution would be the most important elements in a particular signature, and conversely, the influence of the majority of genes became weaker and weaker, or even was over, in that signature when they lay at the center of the distribution [20, 21]. Based on this, oCEM provided the users with three optional post-processing steps attached with ICA and IPCA (two for ICA and one for IPCA) to detect coexpressed gene modules.

For the first option of the post-processing step (“ICA-FDR” assigned to the *method* argument of the function *overlapCEM)*, oCEM did the extraction of non-Gaussian signatures by ICA (the *fastICA* algorithm was configured using *parallel* extraction method and the default measure of non-Gaussianity *logcosh* approximation of negentropy with *α*=1), then the *fdrtool* R tool [31] modeled those signatures as a two-distribution mixture (null and alternative). The null (Gaussian) distribution was fitted around the median of the signature distribution. At last, a user-defined probability threshold (e.g., 0.1, 0.01, 0.001,…), called tail area-based false discovery rate (FDR), was chosen to distribute genes to modules on the condition that a gene whose FDR lesser than the threshold at a signature was assigned to that signature (module). Here we suggested the selection of the sufficiently stringent threshold of 0.001 if appropriate for robustness.

The second option (“ICA-Zscore” assigned to the *method* argument of the function *overlapCEM)* was similar to the first one, but oCEM first did z-score transformation for genes in each signature. A gene belonged to a module if the absolute of its z-score was greater than a user-defined standard deviation threshold (e.g., 0.5 σ,1 σ, 1.5 σ,…). We suggested choosing the sufficiently strict threshold of 3 σ on either side from the zero mean, which picks only out a few genes in the tails of the distributions at any time as possible.

The last option (“IPCA-FDR” assigned to the *method* argument of the function *overlapCEM*) was similar to the first one, but here oCEM used IPCA (the *ipca* algorithm was configured using *deflation* extraction method and the default measure of non-Gaussianity *logcosh* approximation of negentropy with *α*=1) instead of ICA. This algorithm was more robust to noise.

Genes at both extremes of the distribution were considered hub-genes. The Pearson’s correlations of each resulting co-expressed module to each clinical feature of interest were then calculated and reported in R.

### Performance validation of oCEM

#### Gene expression data

We used three example data, human breast cancer [32], mouse metabolic syndrome [33], and Escherichia coli (E.coli) [34], to illustrate the straightforward use of oCEM as well as be convenient for comparing its ability with other tools. In particular, the first case study, downloaded from the cBioPortal for Cancer Genomics [35, 36], was the METABRIC breast cancer cohort in the United Kingdom and Canada. The gene expression data were generated using the Illumina Human v3 microarray for 1,904 samples. The second case study, related to mouse metabolic syndrome (obesity, insulin resistance, and dyslipidemia), was liver gene expressions from 134 female mice including 3600 physiologically relevant genes. The data were employed by the authors of WGCNA [16] to indicate how to use this tool. Finally, the expression values of 4,296 genes from 805 E.coli samples were downloaded from the DREAM5 network inference challenge website.

#### Comparison of oCEM with WGCNA and its improved version iWGCNA

We used the two expression data above to validate the performance of oCEM with WGCNA and an improved version of WGCNA proposed by us [37], temporarily called improved WGCNA (iWGCNA) in this study. For WGCNA, we applied it to the gene expression data using the *blockwiseModules* function (v1.69). All tuning parameters were left as default. For iWGCNA, its improvement was that we added an additional step to the gene clustering process, the determination of the optimal cluster agglomeration method for each particular case. All other tuning parameters were set to their default value, except for the selection of the soft-thresholding value [16].

To compare the power of them, we estimated the pairwise Pearson’s correlation coefficients, *r*, between module eigengenes (MEs, characterized by its first principal component) of resulting modules given by WGCNA (wME) and iWGCNA (iME) versus patterns (i.e., sample components) given by oCEM. This helped us to determine which modules could be missed by WGCNA and iWGCNA. Then, g:Profiler (https://biit.cs.ut.ee/gprofiler/gost) (ver *e102_eg49_p15_7a9b4d6;* accessed on 20 Feb. 2021) [38] verified biological processes and KEGG pathways related to those missed modules. Biological processes and KEGG pathways with adjusted P-values ≤ 0.05 (G:SCS multiple testing correction method [38], two-tailed) were considered to be statistically significant. Importantly, as the g:Profiler database did not support to perform the enrichment analysis from a set of genes in E.coli, we decided to use the Gene Ontology (GO) database [39] for the same task instead.

## Results

### Human breast cancer

In our previous study [37], the breast cancer data were used to detect 31 validated breast-cancer-associated genes, and we then clustered those genes to functional modules using iWGCNA. Here, we revisited the results to be convenient for the comparison. Due to the small number of genes, WGCNA failed to identify any co-expressions across the 1,904 breast cancer patients (the 31 genes were in wM0 or called a gray module), while iWGCNA and oCEM indicated two (iM1 and iM2 respective to turquoise and blue modules) and three modules (oM1, oM2, and oM3), respectively. These implied that the ability of iWGCNA and oCEM was better than WGCNA in the co-expressed gene module identification. Figure 2a indicates that oCEM discovered the three co-expressed modules including a corresponding set of genes of which the overlap was allowed. The correlation analyses of the three identified modules were performed automatically by oCEM (Figure 2b). As a result, oM1 showed a significant negative association with the Nottingham prognostic index only. In particular, oM3 was positively significantly correlated with all three clinical features, including the number of lymph nodes, Nottingham prognostic index, and tumor stages of the breast cancer patients. Besides, oCEM also reported the top 10 hub-genes in each of these modules, including *KMT2C*, *BAP1*, *PTEN*, *NF1*, *RUNX1*, *ZFP36L1*, *CDKN1B*, *BRCA2*, *MAP3K1*, and *PIK3CA* in oM1; *CDH1*, *PIK3R1*, *GATA3*, *CDKN2A*, *TBX3*, *SMAD4*, *KRAS*, *RB1*, *MEN1*, and *RUNX1* in oM2; and *KRAS*, *GPS2*, *SF3B1*, *AGTR2*, *RB1*, *NCOR1*, *SMAD4*, *ERBB3*, *FOXO3*, and *NF1* in oM3.

**Figure 2.**
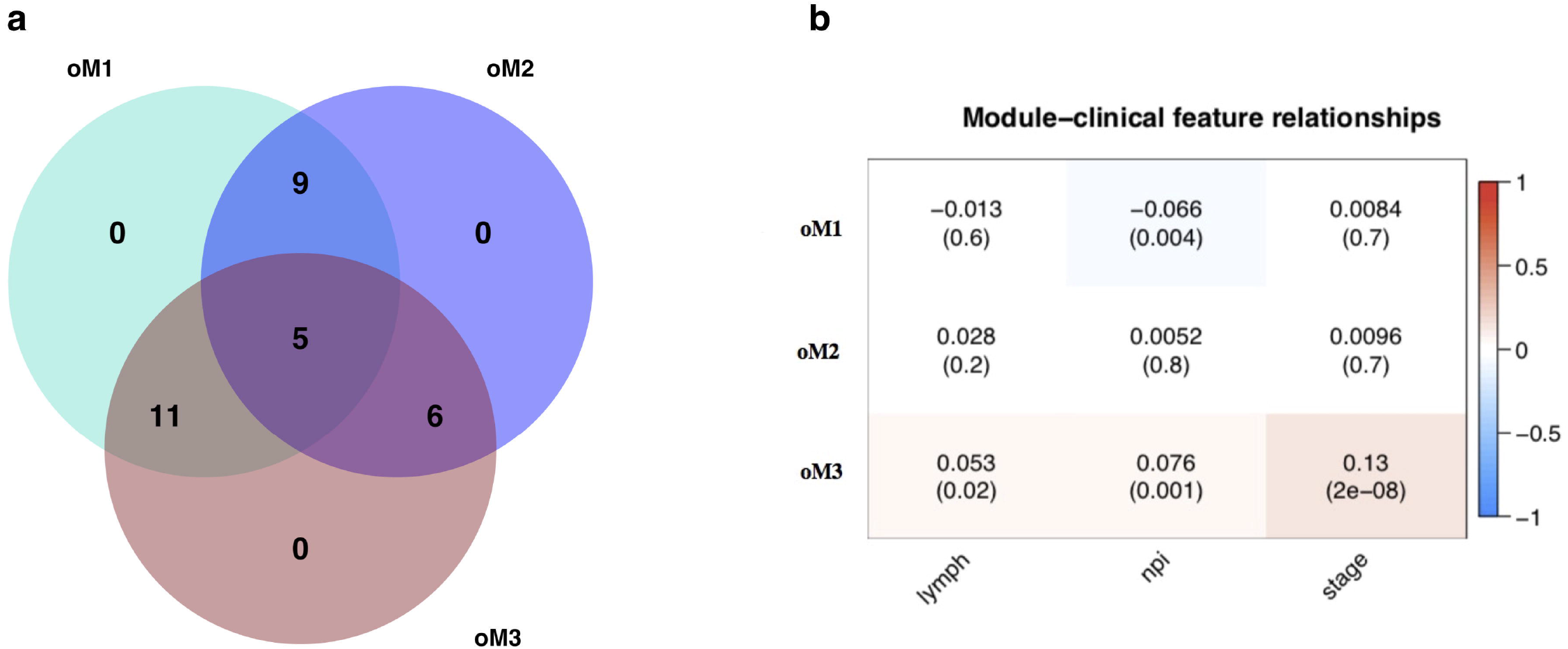
Identification and analysis of functional modules by oCEM. (a) Venn diagram shows the overlap of the 31 driver genes among the three functional modules (oM1, oM2, and oM3). (b) Associations of each module with each of three clinical features of interest. Abbreviation: lymph, the number of lymph nodes; npi, Nottingham prognostic index; stage, tumor stages of all the breast cancer patients; oM, resulting modules generated by oCEM.

We further investigated the power of the three methods by estimating the pairwise Pearson’s correlation coefficients between the one and two modules given by WGCNA and iWGCNA, respectively, versus the three modules given by oCEM as described in the Methods section below. As expected, one wME (100.0%) and one iME (50.0%) respectively showed r > 0.4 with at least one oCEM pattern (i.e., patient components), whereas only 33.3% of oCEM patterns correlated to at least one ME obtained by both WGCNA and iWGCNA with the same intensity (Figure 3a,b and Additional File 2:TableS1). Collectively, both WGCNA and iWGCNA potentially missed two modules oM2 and oM3 (*r* < 0.4). We functionally enriched the two and realized that they possessed an overlapping set of genes significantly associated with regulation of gene expression and development processes and biological pathways related to cancer in general and breast cancer in particular (Additional File 2:TableS2), suggesting that oCEM was most likely to identify biologically relevant modules that were not represented by WGCNA or iWGCNA modules.

**Figure 3.**
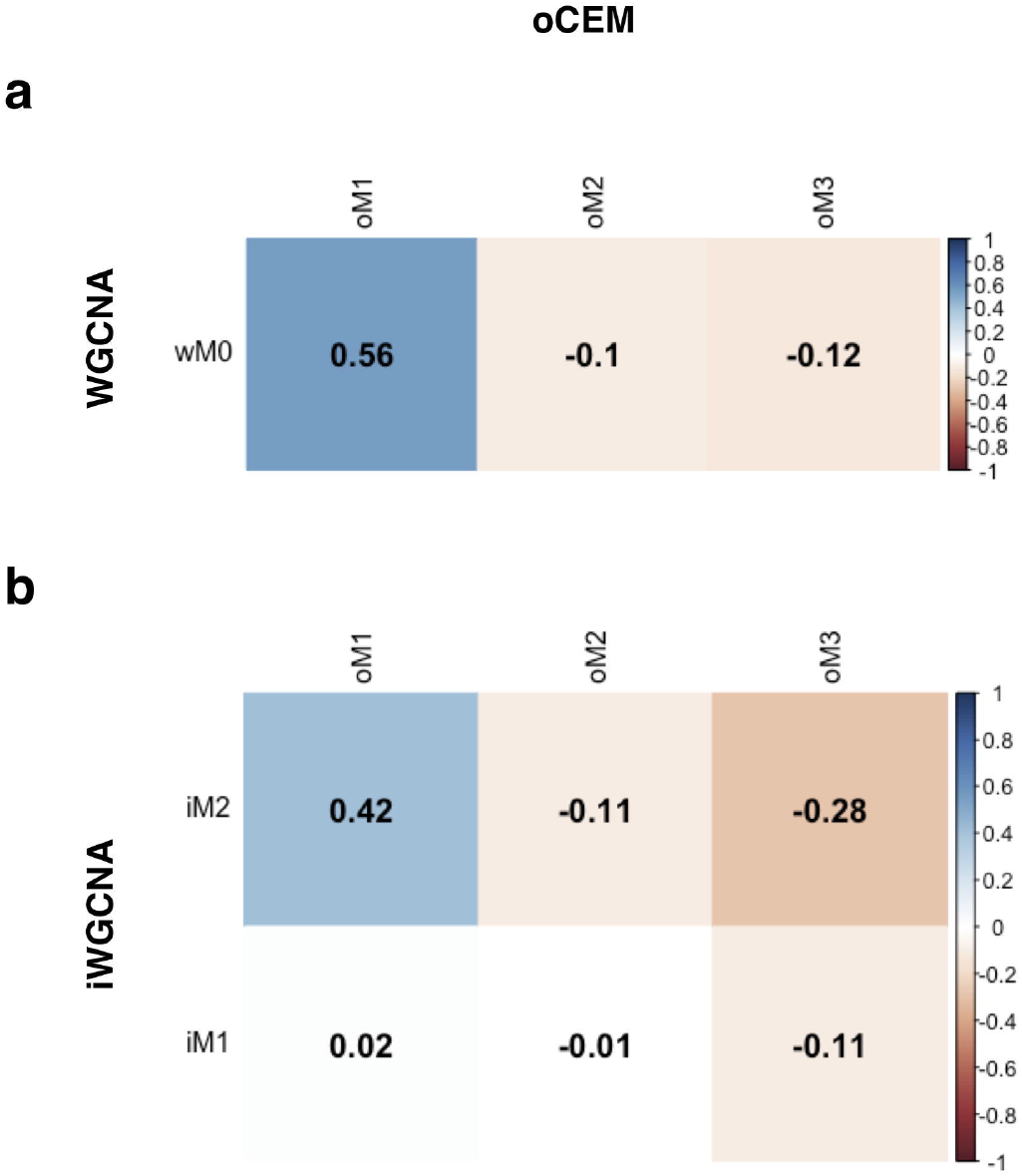
Comparison of identified modules by oCEM with those by WGCNA and iWGNCA. (a) the pairwise Pearson’s correlation coefficients were computed between wM0 versus oM1, oM2, and oM3. (b) the pairwise Pearson’s correlation coefficients were computed between iM1 and iM2 versus oM1, oM2, and oM3. Abbreviation: wM, iM, and oM, resulting modules generated by WGCNA, iWGCNA, and oCEM.

### Mouse metabolic syndrome

Similarly, we applied the three tools to 2,281 gene expressions in the liver of the 134 female mice. As a result, WGCNA, iWGCNA, and oCEM detected 17, 12, and 18 modules, respectively. In this turn, four out of 17 wMEs (23.5%) and four out of 12 iMEs (33.3%) yielded *r* > 0.8 with at least one oCEM pattern (i.e., mouse components). In contrast, those numbers for oCEM were three out of 18 oCEM patterns (16.7%) and four out of 18 oCEM patterns (22.2%) related to at least one wME and one iME with the same intensity, in which WGCNA and iWGCNA could ignore 15 and 14 important oCEM modules (*r* < 0.8), respectively (Additional File 2:TableS3). We analyzed enrichment on those missed modules, rendering all of them associated significantly with relevant metabolic processes and pathways. More details of the preprocessing procedures, analysis processes, and comparisons were shown in Additional File 1.

### E.coli gene expression compendium

Here we did the same as the two studies above once again for 572 gene expressions in a total of 801 samples. To this end, WGCNA, iWGCNA, and oCEM detected 14, nine, and 24 modules, respectively. In this turn, six out of 14 wMEs (42.9%) and five out of nine iMEs (55.6%) issued *r* > 0.8 with at least one oCEM pattern. In contrast, those numbers for oCEM were six out of 24 oCEM patterns (25.0%) and four out of 24 oCEM patterns (16.7%) related to at least one wME and one iME with the same intensity, in which WGCNA and iWGCNA could ignore 18 and 20 important oCEM modules (*r* < 0.8), respectively (Additional File 2:TableS4). We analyzed the biological meaning of the those missed modules, rendering all of them associated significantly with GO terms, which included biological process terms, cellular component terms, and molecular function terms.

## Discussions

Co-expressed gene module identification and sample clustering rely mostly on unsupervised clustering methods, resulting in the development of new tools or new analysis frameworks [16, 37, 40, 41]. However, module detection is unique due to the necessity of ensuring biological reality in the context of gene expressions, such as overlap and local co-expression. In this study, we, therefore, have presented a new tool, oCEM, for module discovery; especially, it differentiates from other advanced methods on the ability to identify different modules which allow having the overlap between them, better reflecting biological reality than methods that stratify genes into separate subgroups. The fact that oCEM outperforms some state-of-the-art tools, such as WGCNA or iWGCNA, in identifying functional modules of genes. Moreover, oCEM is sufficiently flexible to be applied to any organisms, like human, mouse, yeast, and so on. In addition, oCEM is well able to automatically and easily do the two tasks as identification and analysis of modules. These clearly help to support a community of the users with diverse backgrounds, such as biologists, bioinformaticians, and bioinformaticists, who are interested in this field.

When using the decomposition methods, the selection of the optimal number of principal components is vital. Here we also introduce *optimizeCOM* which performs a permutation procedure hoping that the extracted components are generated not-at-random. Based on the three benchmark datasets, including human breast cancer, mouse metabolic syndrome, and E.coli, we can realize that most modules indicated by *optimizeCOM* are highly similar to those displayed by WGCNA and iWGCNA, whereas the rest are new modules significantly associated with clinical features as well as biological processes and pathways. Although further studies are required, these results imply that *optimizeCOM* could provide a suggestion having a high value of reference before using the decomposition methods.

However, we acknowledge that there have still several restrictions of oCEM. Firstly, the input of oCEM is only gene expression matrix. Many prior studies have claimed that integration of multi-omics will enable us to discover molecular mechanisms missed by using each omics technology [40, 42, 43]. For example, as the interpretation of the target expression change is based partly on the change in transcription factor expression [34], this information included may help to improve the ability of module identification. In the future, we will find the way to combine -omics data, possibly including transcription factors, prior to being the input of oCEM. Secondly, each oCEM module consists of many genes, so biologists have difficult experimentally validating such a module by follow-up wet-lab experiments. This is a common problem of existing coexpression identification methods, including oCEM. We suggest that biologists can select some co-expressed modules of interest possessing well-established co-regulation of genes. Another way is choosing some co-expressed modules associated significantly with clinical features of their interest.

## Conclusion

In conclusion, we believe that the oCEM tool may be useful to improve module detection and discover novel biological insights into complex diseases.

## Availability and requirements

Project name: oCEM

Project home page: https://github.com/huynguyen250896/oCEM Operating system(s): Any

Programming language: R Other requirements: None License: MIT

Any restrictions to use by non-academics: none

## Declarations

### Ethics approval and consent to participate

Not applicable

### Consent for publication

Not applicable

### Availability of data and materials

R package of oCEM in the study is available under the MIT license on GitHub at (https://github.com/huynguyen250896/oCEM). Alternatively, the users can also download the raw data of breast cancer from the cBioPortal for Cancer Genomics (http://www.cbioportal.org) [35, 36] under the accession number EGAS00000000083, mouse metabolic sybdrome from the Gene Expression Omnibus (GEO; http://www.ncbi.nlm.nih.gov/geo) under the accession number GSE2814, and E.coli from the DREAM5 network inference challenge website [34] (synapse.org/#!Synapse:syn2787209/wiki/70349) under the identifier syn3130842. Approval by a local ethics committee was not required, all the raw data can be immediately downloaded for research purposes.

### Abbreviations

oCEM: Overlapping CoExpressed gene Module
WGCNA: weighted gene co-expression network analysis
iWGCNA: improved weighted gene co-expression network analysis
ICA: independent component analysis
PCA: principal component analysis
IPCA: independent principal component analysis
E.coli: Escherichia coli
GO: Gene Ontology

### Author contributions

Conception and design of the study: Q-HN; design and implementation of oCEM: Q-HN; computational analysis and result interpretation: Q-HN; manuscript drafting: Q-HN; review and editing: D-HL; supervision: D-HL; The authors read and approved the final manuscript.

## Acknowledgements

Not Applicable

**File name:** Additional File 1.

**File format including the three-letter file extension:** DOC **Title of data:** User manual

**Description of data:** Tutorial and use examples of oCEM.

**File name:** Additional File 2.

**File format including the three-letter file extension:** xlsx **Title of data:** Supporting results.

**Description of data:** Supporting results.

