## Additional File 1 for "oCEM: Automatic detection and analysis of overlapping co-expressed gene modules"

**Additional File 1: Supplementary Materials**

### 1. Supplementary Methods

#### 1.1 ICA and IPCA algorithms

Here we have decided to integrate ICA [1-3] and IPCA [4], assigned to the decomposition methods, into oCEM. Importantly, we have excluded principal component analysis (PCA) [5] method. Figure S1 shows the example output of the decomposition methods, including signatures and patterns with the number of components set to two.

**Figure S1.** Heatmap shows the example output of the ICA and IPCA methods with *k*= 2 (*k*, number of principal components).

#### 1.1.1 ICA algorithm

There are a number of “source signals” (or “original signals” or “statistically independent components”), but due to some external circumstances, only linear mixtures of the source signals are gained. The goal of ICA [1-3] is to recover independent source signals from a mixture of signals. ICA is simply written as a matrix decomposition:

$$X=SA$$

Where $X= x_{mn}$ is normalized data matrix, $S=s_{mk}$ is the impact of component *k* on gene expression level *m* (the first matrix product, vertical rectangle in Figure S1), and $A= a_{kn}$ is expression change of that component in individual *n* (the second matrix product, horizontal rectangle in Figure S1). These *k* components must be as statistically independent as possible.

In the first matrix product, ICA creates nearly sparse signatures, meaning that the majority of their impacts close to 0; nevertheless, each of them possess several genes for which its impacts are considerably different from 0 (genes being at both extremes of the distribution). Mathematically, each of the *k* columns of sub-matrix $S$, considered as “signatures” of the hidden biological processes, is a linear combination of the component on each gene expression (each row of $S$). Especially, different signatures may possess a overlapping set of those genes

In the second matrix product, rows of sub-matrix $A$ represent expression change of the *k* components in individuals. In other words, each pattern reflects the change in expression level in individuals.

#### 1.1.2 IPCA algorithm

The idea of IPCA [4] is a combination of ICA and PCA. Specifically, it first uses PCA in order to reduce the dimension of the data and generate the loading vectors, then applies the fastICA algorithm to those loading vectors, rendering independent principal components (IPC). This makes IPCA more robust to noise.

#### 1.2 optimizeCOM algorithm

After running ICA or IPCA for extracting the components is usually to specify how many components are optimal. The goal is to identify the minimal set of components that can be used to describe the co-expressed gene modules. Importantly, there is not a gold standard for this work. In this study, we present the function *optimizeCOM* in oCEM in the hope of helping users to overcome the problem. The statistical method behind the function *optimizeCOM* is based on random permutations adapted from [6]. *optimizeCOM* first permutes individuals independently for each gene expression in the original data (the certain number of permutations defined by the user, say *P* times) and generates *P* respective random matrices (Figure S2a). Then, it applies the singular value decomposition (SVD) to the original matrix and every resulting random matrices, rendering (1 + *P*) vectors of singular values $\sigma$ corresponding to each of the (1 + *P*) matrices. Suppose we have *k* genes. We then turn those (1 + *P*) singular vectors to the percentage variance explained (%VE) of the total variance as follows:

$${\%VE}_{i,m}=\frac{\sigma_{i,m}}{\sum_{i=1}^{k} {\sigma_{i,m}}^{2}} \times100 (\%), (m=\bar{1,1+P})$$

where $\sigma_{i,m}$ is the *k* singular values in the *m*^th^ matrix, which can be interpreted as the correlation of each gene with that matrix. $\sum_{i=1}^{k} {\sigma_{i,m}}^{2}$ are the sum of squared singular values across the *k* genes for each matrix, which represents the total amount of variance that can be explained by the matrix.

Next, average (over *P* random matrices) the first %VE, the second %VE, …, the *P*^th^ %VE. At last, the (first) intersection between a curve obtained from %VE of the parental matrix and a curve obtained from the mean %VE of all the *P* random matrices plotted in the Screeplot is the optimal component number, considering that beyond this number, components mostly reflected noise (Figure S2b). If the intersection point is a decimal number on the horizontal axis, *optimizeCOM* rounds up to the next number; e.g., 8.2 or 8.8 turn to 9.

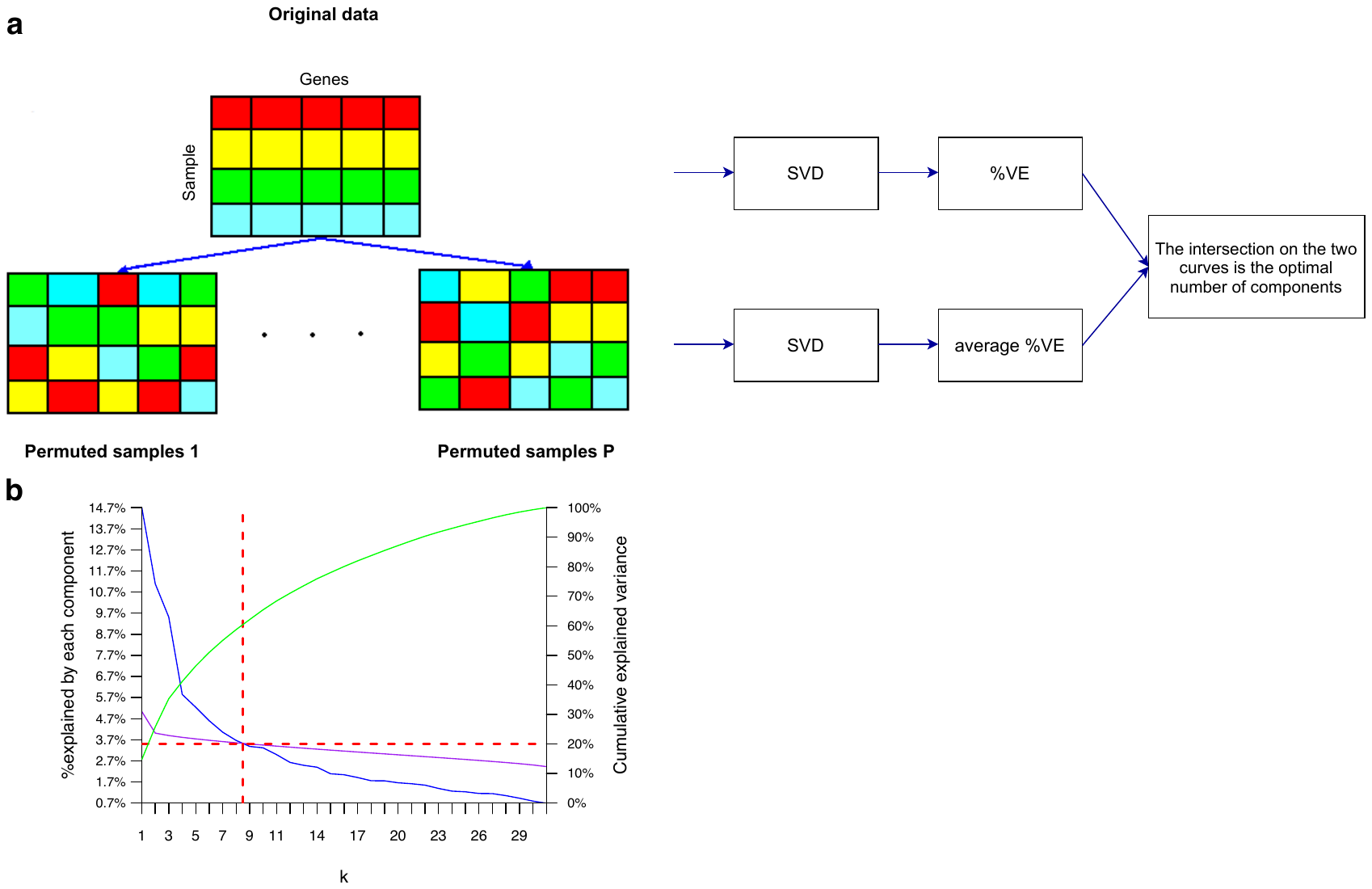

**Figure S2.** (a) pipeline of the permutation procedure of *optimizeCOM* where all the samples (rows) are permuted independently for each gene (each column). It then applies the singular value decomposition (SVD) to both the original data and every random matrices, and infers the optimal number of components based on the intersection between the two curves created by the percentage of variance explained (%VE) of the real data and average %VE of all the *P* random matrices. (b) The example screeplot illustrates %VE (left Y-axis) and the culmulative VE (right Y-axis, calculated based on %VE) by the principal components *k* (X-axis) of the SVD. The blue solid curve shows %VE and the green curve shows the cumulative VE of the real data matrix across principal components. The purple solid curve presents average %VE of the *P* random matrices across principal components. The optimal number of components is determined at the intersection between the blue curve (observed variance) and the purple curve (variance expected under random). The optimal number is approximately 9 (red vertical line).

It is crucial to consider how big *P* needs to be as Horn *et al* [6] have also mentioned this issue. Moreover, simulation studies have reported that *P* < 99 have low power, whereas *P* $\geq$ 499 is recommended [7-9]. Additionally, *P* = 1000 has been proved to produce good results [9, 10]. From that, 1000 is left as default in *optimizeCOM*. The user should be aware that an increase in numbers of permutations will lead to a significant increase in computation time with possibly only a small gain on the approximation.

Also, to suggest which method (ICA or IPCA) should be selected, *optimizeCOM* runs idenpendently the two methods for the input data matrix and check whether all principal components generated from each method have all their kurtosis values < 3 or not. If a certain method issues all principal components whose kurtosis < 3, it should be ignored.

### 2. Supplementary Results

oCEM is an R package can be freely available on our Github repository ([https://github.com/huynguyen250896/oCEM](https://github.com/huynguyen250896/overlappingCGM)). To show a straightforward use of oCEM, we re-use -omic data used in our prior study [11], downloaded from our github repository ([https://github.com/huynguyen250896/oCEM/tree/main/data_n_code/human%20breast%20cancer](https://github.com/huynguyen250896/overlappingCGM/tree/main/data_n_code/human%20breast%20cancer)) or the cBioPortal for Cancer Genomics (<http://www.cbioportal.org)> [12, 13], including gene expression (EXP; n = 1,904), in a cohort of breast cancer patients.	Besides, we also apply our tool to mouse metabolic syndrome containing liver EXP from female mice (n =135) [14]. The raw data and R codes for pre-processing processes can be seen in ([https://github.com/huynguyen250896/oCEM/tree/main/data_n_code/mouse%20metabolic%20syndrome](https://github.com/huynguyen250896/overlappingCGM/tree/main/data_n_code/mouse%20metabolic%20syndrome)). Finally, we apply our tool to E.coli expression data, including 4,296 genes from 805 E.coli samples, downloaded from the DREAM5 network inference challenge website [15]. The raw data and R codes for pre-processing processes can be seen in (<https://github.com/huynguyen250896/oCEM/tree/main/data_n_code/ecoli>).

#### 2.1 human breast cancer

In our previous study [11], we have discovered 31 driver genes associated with breast cancer. Here we first preprocess the gene expression matrix (exp) following the requirement of oCEM, including its rows are the 1904 patients and its columns are those 31 genes. Note that normalization does not need to be done in advance because oCEM can do this task on its own. Also, we must preprocess the clinical data (clinicalEXP), including the selection of clinical features of interest, and turning its data structure into required one: its rows are the 1904 samples and its columns are clinical features of those patients. In the present study, we re-select three clinical features: the number of lymph (lymph), Nottingham prognostic index (npi), and tumor stages (stage) selected in the previous work already. Importantly, before we establish co-expressed gene modules, we need to exclude outliers in the exp data. However, since we do not see any outlier in the breast cancer exp data, we do not show R codes below. The user can see the R codes for this task in the second case, i.e., mouse metabolic syndrome.

Load necessary libraries

```
# set up the working directory
setwd("~/path/to/your/raw/data")

# library
devtools::install_github("huynguyen250896/oCEM", force = T) #NOTE: Download all dependencies of the tool!

x=c("oCEM", "dplyr", "dynamicTreeCut", "flashClust","Hmisc", "WGCNA", "moments", "fastICA", "tidyr", "fdrtool", "mixOmics", "cluster", "purrr")
lapply(x, require, character.only = TRUE)

# load raw data
exp = read.table('data_mRNA_median_Zscores.txt', sep = '\t', check.names = FALSE, header = TRUE, row.names = NULL)

clinical = read.table('data_clinical_patient.txt', sep = '\t', check.names = FALSE, header = TRUE, row.names = 1,fill=TRUE)

# 31 identified driver genes
driver=c("MAP2K4", "ARID1A", "PIK3CA", "TBX3", "MAP3K1", "TP53", "AKT1", "GATA3", "CDH1", "RB1", "CDKN1B", "NCOR1", "CDKN2A", "ERBB2", "KRAS", "BRCA2", "BAP1", "PTEN", "CBFB", "KMT2C", "RUNX1", "NF1", "PIK3R1", "ERBB3", "FOXO3", "SMAD4", "GPS2", "AGTR2", "ZFP36L1", "MEN1","SF3B1")
length(driver) #31 driver genes

# only keep the 31 driver genes in exp
exp=exp %>%
  dplyr::filter(.$Hugo_Symbol %in% driver) %>%
  tibble::column_to_rownames('Hugo_Symbol') %>%
  dplyr::select(-Entrez_Gene_Id)

# check dimension
dim(exp) # 31 1904
dim(clinical) # 2509   21

# match patients sharing between exp versus clinical
# exp are the matrix whose rows are samples and columns are genes
exp = exp[,intersect(colnames(exp), rownames(clinical))] %>% t()
clinicalEXP = clinical[intersect(rownames(exp), rownames(clinical)), ]

# preprocess clinicalEXP
clinicalEXP = clinicalEXP %>%
  tibble::rownames_to_column('sample') %>%
  dplyr::select(c(sample, LYMPH_NODES_EXAMINED_POSITIVE, NPI, stage)) %>%
  tibble::column_to_rownames('sample')

colnames(clinicalEXP)[1:3] =  c("lymph", "npi", "stage")
str(clinicalEXP)
# 'data.frame':	1904 obs. of  5 variables:
# $ lymph : int  1 5 8 1 0 1 0 2 0 6 ...
# $ npi   : num  4.04 6.03 6.03 5.04 3.05 ...
# $ stage : int  2 2 3 2 2 2 1 2 2 4 ...
```

We apply WGCNA to the data with all tuning parameters are left as default, except for the minimum module size at 10 due to having the 31 genes only.

```
# WGCNA
wgcna = blockwiseModules(exp, power = 6,
    TOMType = "unsigned", minModuleSize = 10,
    reassignThreshold = 0, mergeCutHeight = 0.25,
    numericLabels = TRUE, pamRespectsDendro = FALSE,
    verbose = 3)
```

For oCEM, we first use optimizeCOM to determine the optimal number of principal components and which method should be selected. Then, optimizeCOM suggests choosing nine components and the ICA method. We then input the expression data exp and clinical data clinicalEXP to oCEM with those suggested selections. Here we feed ICA-Zscore to the method argument for illustrative purposes as this post-processing step does not create oM0 (typically called a gray module that include genes not co-expressed with one another).

```
# oCEM
optimizeCOM(data = exp)
# >> oCEM suggests choosing the optimal number of components is: 9
# >> oCEM also suggests using ICA for your case.

cem=overlapCEM(data = exp, clinical = clinicalEXP, ncomp = 9,
                        method = 'ICA-Zscore', cex.text = 1.0)
```

For iWGCNA, we use R code released in our Github (<https://github.com/hauldhut/drivergene>) [11].

```
# The following setting is important, do not omit.
options(stringsAsFactors = FALSE);
enableWGCNAThreads() ### Allowing parallel execution

# we choose the soft-thresholding of 6 based on our prior work
softPower = 6;
adjacency = adjacency(exp, power = softPower,
                      type = "signed");

# Turn adjacency into topological overlap
TOM = TOMsimilarity(adjacency, TOMType = "signed");
dissTOM = 1-TOM

# Hierichical clustering
# Seek the optimal agglomeration method
# methods to assess
m <- c( "average", "single", "complete", "ward")
names(m) <- c( "average", "single", "complete", "ward")
# function to compute agglomerative coefficient
set.seed(25896)
ac <- function(x) {
  agnes(exp, method = x)$ac
}

map_dbl(m, ac) # Agglomerative coefficient of each agglomeration method
# average    single  complete      ward
# 0.4136737 0.3510251 0.4815638 0.5563910

# assign gene names from adjacency to dissTOM
rownames(dissTOM) = rownames(adjacency)
colnames(dissTOM) = colnames(adjacency)
# Call the hierarchical clustering function
geneTree = hclust(as.dist(dissTOM), method = "ward.D2");

# We set the minimum module size at 10:
minModuleSize = 10;

# Module identification using dynamic tree cut:
dynamicMods = cutreeDynamic(dendro = geneTree, distM = dissTOM,
                            minClusterSize = minModuleSize);
table(dynamicMods)
# dynamicMods
# 1  2
# 16 15

# Convert numeric lables into colors
moduleColors = labels2colors(dynamicMods)

# Define numbers of genes and samples
nGenes = ncol(exp);
nSamples = nrow(exp);
# Recalculate MEs with color labels
MEs0 = moduleEigengenes(exp, moduleColors)$eigengenes
MEs = orderMEs(MEs0)
```

At last, we compare the performance of oCEM with that of WGCNA and iWGCNA as follows:

```
# oCEM versus WGCNA
cor(cem$patterns, wgcna$MEs)

# oCEM versus iWGCNA
cor(cem$patterns, MEs)
```

The comparative results are shown in TableS1 in Additional File 2.

#### 2.2 mouse metabolic syndrome

Firstly, we load the raw data including EXP and its clinical data (exp and cli)

```
# load raw file
exp = read.table("LiverFemale3600.csv", header = T, check.names = F, sep=",")
cli = read.table("ClinicalTraits.csv", header = T, check.names = F, sep=",", row.names = 1)
```

We remove unknown genes (i.e., coded as 0 in the data) and duplicated genes due to insufficient information to retain them.

```
#remove missing gene names and duplicated genes in exp
exp = exp[which(exp$gene_symbol != "0"),]

dup = duplicated(exp$gene_symbol)
exp = exp[which(dup == FALSE),]

# turn exp1 into satisfactory format
# requires data whose rows are samples and columns are genes.
exp = exp %>%
  dplyr::select(-c(substanceBXH, LocusLinkID, ProteomeID, cytogeneticLoc,
            CHROMOSOME, StartPosition, EndPosition)) %>%
  tibble::remove_rownames() %>%
  tibble::column_to_rownames('gene_symbol') %>%
  drop_na() %>% t()
```

Since oCEM is sensitive to outlier samples, we find potential ones by using the hierarchical clustering method. Finally, we exclude the mouse named F2_221 (Figure S3).

```
# detect outliers
sampleTree = hclust(dist(exp), method = "average")
par(cex = 0.6);par(mar = c(0,4,2,0))
plot(sampleTree, main = "Sample clustering to detect outliers", sub="", xlab="",
     cex.lab = 1.5,cex.axis = 1.5, cex.main = 2)

# Plot a line to show the cut
abline(h = 12.2, col = "red");
# Determine cluster under the line
clust = cutreeStatic(sampleTree, cutHeight = 12.2, minSize = 10)
table(clust)
# clust 1 contains the samples we want to keep.
keepSamples = (clust==1)
exp = exp[keepSamples, ]
```

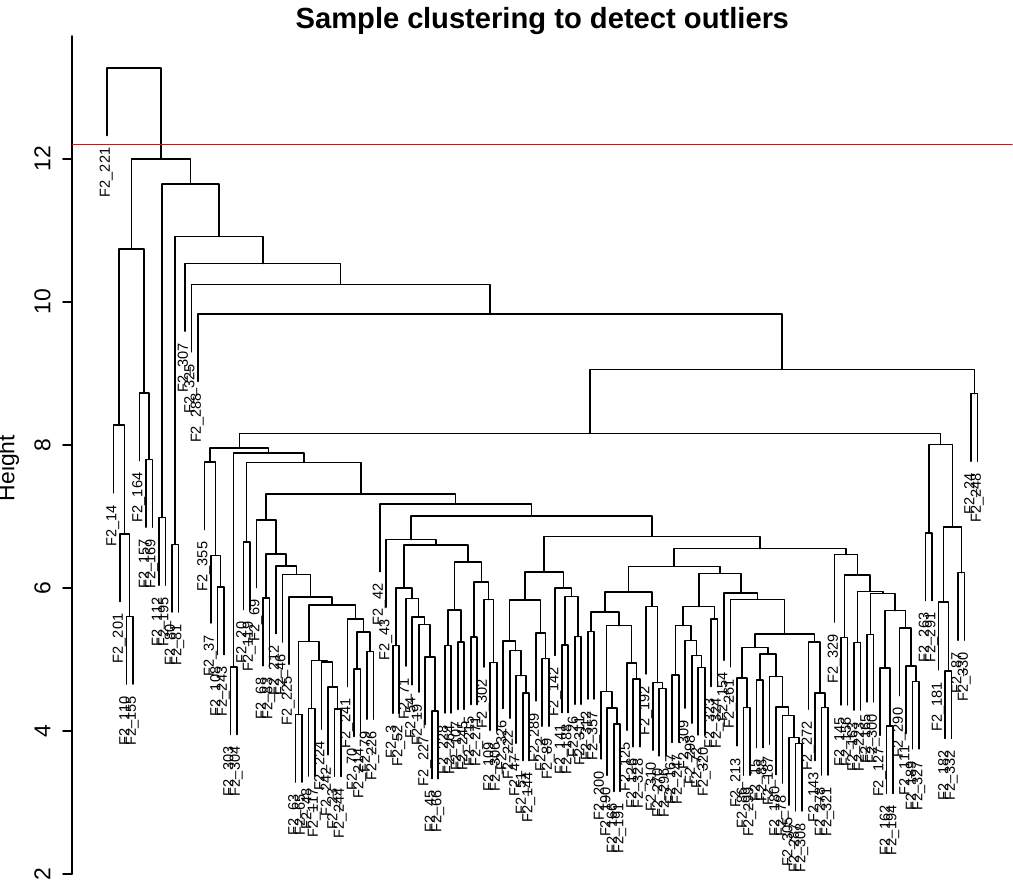

**Figure S3.** Detect and remove outliers.

We keep the mice that share between exp and cli at their rows and in exactly the same order. We also only keep eight out of 20 clinical features; including body weight weight_g, body length length_cm, abdominal fat ab_fat, total fat total_fat, ulcerative colitis UC, free fatty acids FFA, glycemic index Glucose, two LDL and VLDL cholesterol levels LDL_plus_VLDL (Figure S4).

```
# match mouses that share between cli versus exp
cli = cli[cli$Mice %in% rownames(exp),]
cli = cli %>%
  remove_rownames() %>%
  tibble::column_to_rownames('Mice') %>%
  dplyr::select(-c(Number, sex, Mouse_ID, Strain, DOB, parents, Western_Diet,
            Sac_Date, comments, Note))

# how the clinical traits relate to the sample dendrogram.
# Re-cluster samples
sampleTree2 = hclust(dist(exp), method = "average")
# Convert traits to a color representation: white means low, red means high, grey means missing entry
traitColors = numbers2colors(cli, signed = FALSE);
# Plot the sample dendrogram and the colors underneath.
plotDendroAndColors(sampleTree2, traitColors,groupLabels = names(cli), main = "Sample dendrogram and trait heatmap")
# white means a low value, red a high value, and grey a missing entry.
```

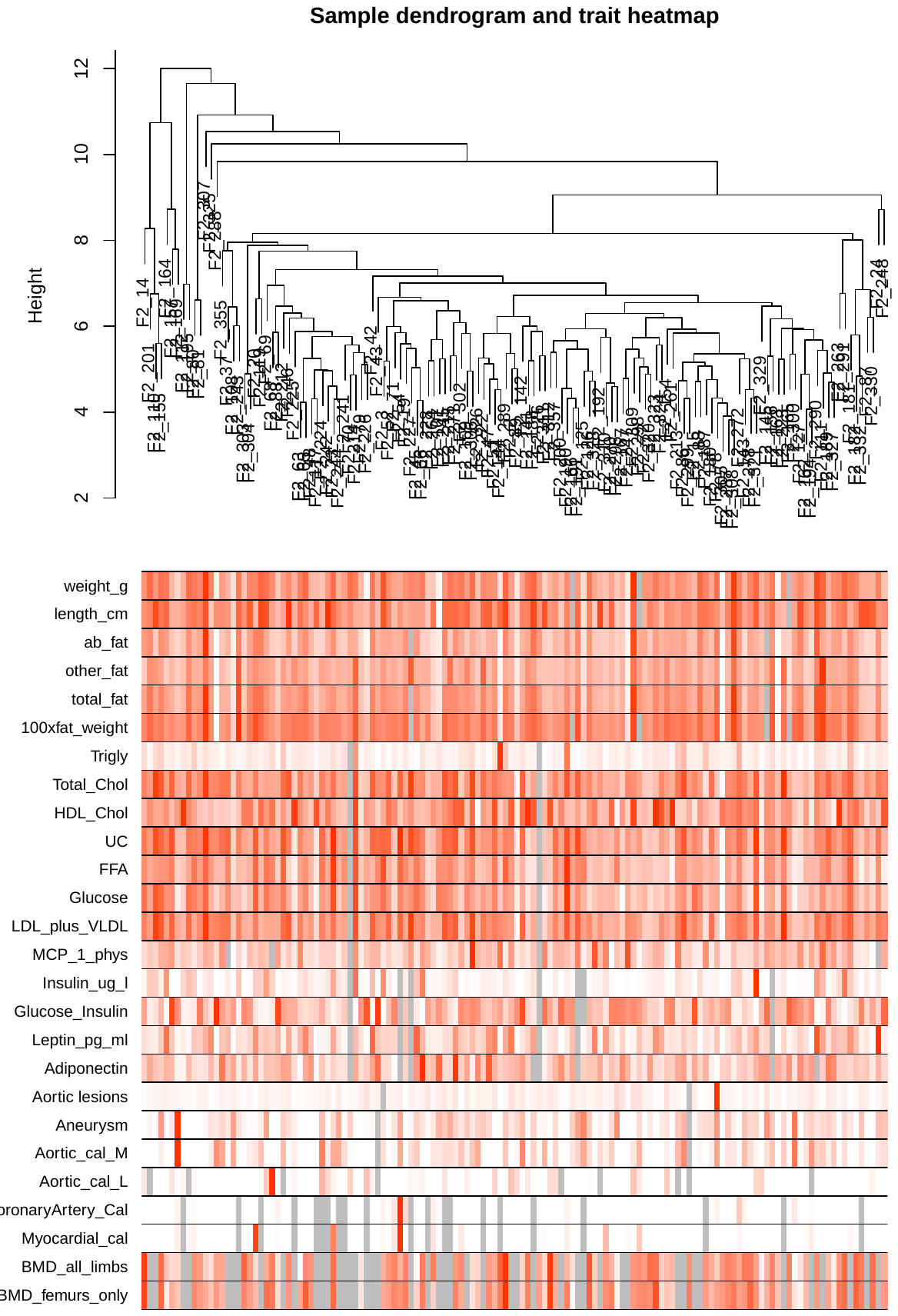

**Figure S4.** How the clinical features relate to the sample dendrogram. White means a low value, red a high value, and grey a missing entry

```
# Only keep several clinical features with red color
cli = cli %>%
  dplyr::select(weight_g, length_cm, ab_fat,
                total_fat, UC, FFA, Glucose,
                LDL_plus_VLDL)

# make sure that mice that share between exp and cli are included at their rows and in exactly the same order
all(rownames(exp) == rownames(cli))
#[1] FALSE
exp = exp[rownames(cli), ]

# check dimension
dim(exp)
#[1] 134 2281
dim(cli)
#[1] 134  8
```

After the preprocessing step, we again apply WGCNA to the data with defaut parameters.

```
wgcna = blockwiseModules(exp, power = 6,
        TOMType = "unsigned", minModuleSize = 30,
        reassignThreshold = 0, mergeCutHeight = 0.25,
        numericLabels = TRUE, pamRespectsDendro = FALSE,
        verbose = 3)
```

Similarly, the following are R codes for applying oCEM to the data. In this turn, optimizeCOM indicates that 18 components are the best but cannot indicate which method should be selected. Finally, we realize that ICA-Zscore for illustrative purposes.

```
# oCEM
optimizeCOM(data = exp)
# >> oCEM suggests choosing the optimal number of components is: 18
# >> Both ICA and IPCA-FDR are appropriate for your case. Please use another more stringent approach to make the best decision.

cem=overlapCEM(data = exp, clinical = cli, ncomp = 18,
                        method = 'ICA-Zscore')
```

And for iWGCNA

```
# The following setting is important, do not omit.
options(stringsAsFactors = FALSE);
enableWGCNAThreads() ### Allowing parallel execution

#we realize that the soft-thresholding of 5 is best fit for the data
softPower = 5;
adjacency = adjacency(exp, power = softPower,
                      type = "signed");

# Turn adjacency into topological overlap
TOM = TOMsimilarity(adjacency, TOMType = "signed");
dissTOM = 1-TOM

# Hierichical clustering
# Seek the optimal agglomeration method
# methods to assess
m <- c( "average", "single", "complete", "ward")
names(m) <- c( "average", "single", "complete", "ward")
# function to compute agglomerative coefficient
set.seed(25896)
ac <- function(x) {
  agnes(exp, method = x)$ac
}

map_dbl(m, ac) # Agglomerative coefficient of each agglomeration method
# average    single  complete      ward
# 0.4136737 0.3510251 0.4815638 0.5563910

#assign gene names from adjacency to dissTOM
rownames(dissTOM) = rownames(adjacency)
colnames(dissTOM) = colnames(adjacency)
# Call the hierarchical clustering function
geneTree = hclust(as.dist(dissTOM), method = "ward.D2");

# We set the minimum module size at 10:
minModuleSize = 10;

# Module identification using dynamic tree cut:
dynamicMods = cutreeDynamic(dendro = geneTree, distM = dissTOM,
                            minClusterSize = minModuleSize);
table(dynamicMods)
# dynamicMods
# 1   2   3   4   5   6   7   8   9  10  11  12
# 341 311 272 230 216 211 188 168 112  84  80  68

# Convert numeric lables into colors
moduleColors = labels2colors(dynamicMods)

# Define numbers of genes and samples
nGenes = ncol(exp);
nSamples = nrow(exp);
# Recalculate MEs with color labels
MEs0 = moduleEigengenes(exp, moduleColors)$eigengenes
MEs = orderMEs(MEs0)
```

At last, we compare the performance of oCEM with that of WGCNA and iWGCNA as follows:

```
#oCEM versus WGCNA
cor(cem$patterns, wgcna$MEs)

#oCEM versus iWGCNA
cor(cem$patterns, MEs)
```

The comparative results are shown in TableS3 in Additional File 2.

### 3. Understanding the tool and its results

#### 3.1 optimizeCOM function

optimizeCOM(data = NULL, P=1000, standardize = T, verbose = T)

where the argument data is gene expression matrix whose rows are samples and columns are genes. P is the number of permutations which is set to 1000 as default as explained above. If your expression matrix is not standardized, please feed T/TRUE to the argument standardize to let oCEM do this.

When you correctly input your data and the parameters to optimizeCOM,	it will automatically output in the R console result as follows (Figure S5).

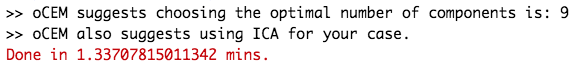

**Figure S5.** optimizeCOM runs with the input of the 31 driver genes in human breast cancer and P = 1000. The results are printed out in the R console result.

#### 3.2 overlapCEM function

overlapCEM(data = NULL, clinical = NULL, ncomp = NULL, standardize = T, method = c("ICA-FDR", "ICA-Zscore", "IPCA-FDR"), cex.text = NULL)

where the argument data is gene expression matrix whose rows are samples and columns are genes. clinical is clinical data including clinical features of choice. If your expression matrix is not standardized, please feed T/TRUE to the argument standardize to let oCEM do this. The argument ncomp is user-decided number of components that can be refer to the suggestion of optimizeCOM. The argument method shows options which are in a format as “method-post-processing”, in which the methods may be ICA or IPCA and the post-processing procedures may be FDR or Z-score. The argument cex.text helps the user change the font size of texts in cells of resulting Figures generated by oCEM (i.e., Figure S7).

When you correctly input your data and the parameters to oCEM, it will automatically output in the R console result as follows (Figure S6 and Figure S7).

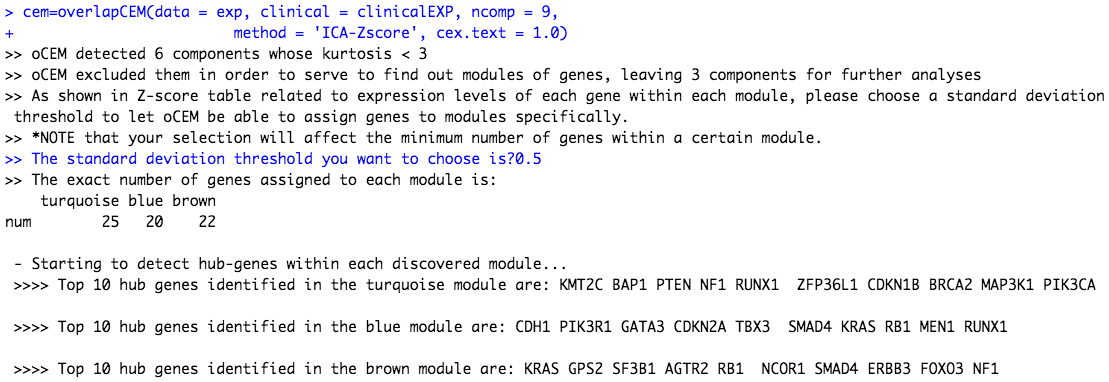

**Figure S6.** oCEM runs with the input of the 31 driver genes in human breast cancer. The results are printed out in the R console result. Firstly, oCEM seeks and removes signatures whose kurtosis value < 3. Then, it asks the user to choose the threshold to distribute genes to modules. Due to the small number of genes, we choose the threshold of 0.5 sigma on either side from the zero mean. Finally, oCEM prints all the results in the R console result.

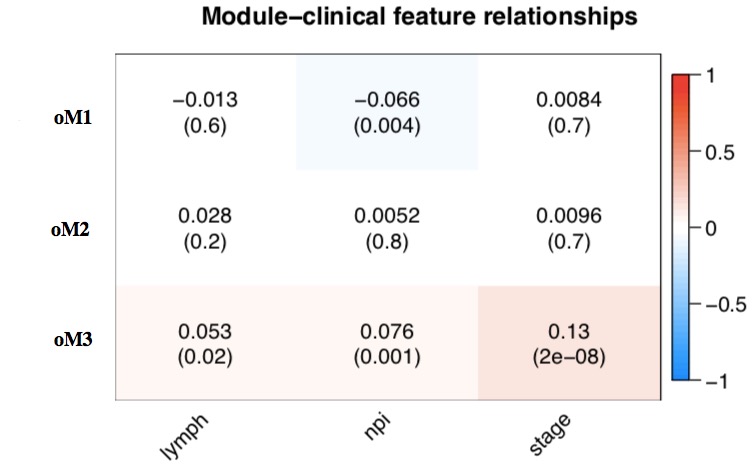

**Figure S7.** Associations between the three resulting modules (oM1, oM2, and oM3) and the clinical features of interest (i.e., lymph, npi, and stage) calculated by the Pearson’s correlation. Abbreviation: the number of lymph node, lymph; Nottingham prognostic index, npi; tumor stages, stage.

Note that the user can specifically see which genes are distributed to which module by the command (Figure S8)

cem$signatures

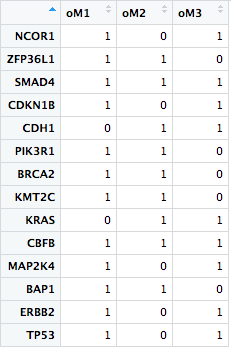

**Figure S8.** Results of which genes (rows) are distributed to which module (columns). 1 = module-assigned, 0 is otherwise.
